# Comparison of Conventional Cytomorphology, Automated Cell Counting, and FTIR Spectroscopy for Detection of CNS Involvement in Acute Leukaemia

**DOI:** 10.1101/2024.07.01.601535

**Authors:** lasaki abiodun esohe, Samson ADEGBE

## Abstract

**Background:** The incidence of central nervous system (CNS) involvement in pediatric leukemia patients presents significant diagnostic challenges. This study compares the diagnostic capabilities of Fourier Transform Infrared (FTIR) Spectroscopy with conventional cytomorphology and automated cell counting to detect CNS involvement in pediatric leukemia.

**Methods:** A cohort of pediatric patients diagnosed with leukemia was assessed at Altınbaş University Medical Park Hospital. Cerebrospinal fluid (CSF) samples were analyzed using FTIR spectroscopy and compared with outcomes from conventional cytomorphology and automated cell counting to evaluate diagnostic efficacy and accuracy.

**Results:** FTIR spectroscopy demonstrated a higher sensitivity in detecting biochemical markers of CNS involvement than conventional methods. Distinct spectral profiles enabled the identification of specific biochemical changes associated with CNS leukemia.

**Conclusions:** FTIR spectroscopy offers a promising alternative to traditional diagnostic methods, providing rapid, non-invasive, and accurate detection of CNS involvement, which is crucial for timely and effective treatment of pediatric leukemia.

## 1. Introduction

Leukemia is the most common type of cancer in children, with CNS involvement indicating a more aggressive and potentially lethal disease course. The early detection of CNS involvement is critical for effective treatment planning. This study evaluates the potential of FTIR spectroscopy, a less invasive and possibly more sensitive technique, to identify CNS involvement early in the disease process compared to traditional methods such as cytomorphology and automated cell counting. Acute lymphocytic leukemia (ALL) and acute lymphocytic leukemia (AML) are the two most common childhood cancers in the world. White blood cells do not develop properly in leukemia patients and stem cells turn into lymphoblasts (leukemia cells) [Samantha et al., 2017]. Anemia, weakness, coldness, pale complexion, and shortness of breath are among the signs and symptoms of childhood leukemia. Infections, fever, and decreased immunity can be caused by a low white blood cell count. It can be an indication of childhood leukemia if an adolescent develops a long-standing illness or has repeated infections. [Samantha et al., 2017]. In leukemia-related children, the involvement of the central nervous system was initially rare (3–5%), but after recurrence it increased considerably, with about 3040 % of these patients presenting with CNS disease (Frishman-Levy, 2017). Progress in cancer therapy, especially in the testicles and extramedullary areas, has reduced non-brain recurrence. It is believed that blood-brain barrier, blood-leptomeningeal barrier and blood-cSF barrier of all endothelial origin affect the movement of leukemia cells into the CNS. According to a mouse study [Yao et al., 2018].

Using cytological examination and magnetic resonance imaging (MRI), central nervous system (CNS) leukemia could be leptomeningeal and visualized in the cerebrospinal fluid [Clark et al., 2010]. The following are the most used diagnostic modalities for acute CSF leukemia in clinical practice: magnetic resonance imaging, cytopathology, and flow cytometry assessment. The additional examination instruments are molecular biological methods, Fourier transform infrared spectroscopy (FTIR) and humoral parameters in the cerebrospinal fluid. With MR imaging, it is often difficult to distinguish the distributed lesions in the brain due to CNS leukemia from CNS infections or neurodegenerative diseases, as they are also common in patients with acute leukemia (Deak et al., 2021). Furthermore, MRI can detect acute CNS leukemia in parenchymal regions of the brain and spinal cord. In recent years, heptomeningeal disease has become a more common diagnosis. In the late 19th century, the first description of cancer patients’ leptomeningeal metastases was published. Childhood leukemia. Cytometric tests are considered gold standards for identifying CSF cancer cells. In 1904, the benefits of cytology in the detection of cancer cells in CSF were documented for the first time. Many subsequent articles corroborated this conclusion. CSF analysis quickly became popular to detect involvement of CNS in hematological diseases, which is a common consequence. Since there was no alternative test that could reliably detect neoplastic involvement in the 1950s, cytomorphology has acquired a new importance. Since 41 percent of the false negative cases were recorded, subsequent autopsy studies have shown that cytology lacks the sensitivity required to detect most of these cases. Because cells have a short lifespan and rapidly degenerate, accurate morphological diagnosis depends on well-preserved cell samples. If cells are not properly treated, they may no longer be useful for diagnosis. Even morphology occasionally results in indistinguishability between benign and malignant lymphocytes, which may explain why cytology is so sensitive to diagnosis. According to Gupta et al. Flowcytometry (FC) is widely used to detect subtypes, diagnose ALL, and monitor minimal residual diseases (MRD). (2018). Recent new definitions of complete remission in all children, based on morphological and FC analysis [Crespo-Solis et al., 2012], have been proposed. suggesting that FC could replace or at least complement the current morphological assessment in ALL (Thastrup et al., 2022). On the other hand, the clinical significance of a positive FC-CSF study has not yet been clearly clarified. However, conventional cytomorphology analysis (CCC) by Cytospin after CSF cytocentrifugation is the gold standard for identifying lymphoblasts in leukemia patients [Glantz et al., 1998]. According to Pui et al. 2009: This approach has about 50% of the sensitivity of the detection of CNS disorders, but more than 95% of the specificity of CNS infiltration. Recent studies have shown that most CNS relapses occur in patients who had not been involved in CNS at the time of CCC diagnosis [de Graaf et al, 2011]. Fluctuation-cytometric immunophenotyping (FCI) is more sensitive than CCC analysis of CSF, and has been shown in many studies to be an effective method for identifying abnormal cell characteristics. In polymerase chain reactions (PCR) studies, CSF samples from ALL pediatric patients were analyzed. The authors found that DNA was best preserved by CSF in RPMI tissue culture mediums without serum in a 1:1 ratio (Ikonomidou, 2021). After induction treatment, the samples were tested for negative PCR. The authors conclude that CSF real-time PCR analysis is a reliable tool for determining CNS leukemia and could be used to improve CNS status classification in the future. The basis of various spectroscopy-based diagnostic tools is that tissue biochemistry changes reflect changes in spectral frequencies. Among the rapidly developing technologies is Fourier Transformed Infrared Spectroscopy (FTIR), which will enable a faster, faster, and more objective diagnosis in the coming decade (Diem et al., 2002). Over the past decade, FTIR spectroscopy has become a useful tool for medicine. Enhanced FTIR microscopy is enabling objective diagnosis (Diem and others, 1999; Ramesh et. al., 2001).

Monitoring cellular changes using FTIR spectroscopy is also an effective and non-destructive approach [Banyay et al, 2003]. Several studies have been published emphasizing the utility of spectroscopic approaches in cancer detection [Mostaco-Guidolin et al., 2009]. However, most focus on characterizing and differentiating cells and tissues by assessing a collection of bands rather than each band individually. In recent years, FTIR spectroscopy and micro spectroscopy have been used for early detection of various types of cancer by detecting chemical changes in tissue, cell and body fluid samples (Chaber et al., 2021). The General Information section of Dissertation 2 describes in detail the use of FTIR and micro spectroscopy for early detection of cancer. Leukemia could be identified by FTIR based on the lipid and DNA concentration and the general spectrum properties [Schultz et al., 1996]. However, not many studies have been conducted on the use of FTIR spectroscopy to detect cancer in children. It is unclear whether FTIR spectroscopy is useful as a diagnostic or prognostic tool in leukemia as it has not been adequately studied in this disease.

The investigation is motivated by the need for more accurate and timely diagnostic techniques that can influence treatment outcomes positively. By integrating advanced spectroscopic analysis, this study aims to refine the diagnostic approach for CNS involvement in leukemia, potentially setting new standards for clinical practice.

## 2. Methods

This study was conducted using a cross-sectional design where CSF samples from pediatric leukemia patients were analyzed. Samples were collected under stringent conditions to maintain integrity and were subsequently analyzed using three diagnostic approaches: conventional cytomorphology, automated cell counting, and FTIR spectroscopy. The latter employs advanced spectral analysis techniques to detect unique molecular signatures indicative of CNS involvement.

### HUMAN STUDIES

Human cerebrospinal fluid (CSF) samples were used for this study. CSF samples were collected from Altinbas University Medical Park Hospital from the pediatrics department from children within the age of 4-7 years in Istanbul Turkey. Informed consent was obtained from the legal guardians of all patients before the procedure. However, samples used for this work were patients with Acute Lymphocytic Leukemaia, the procedure is a routine procedure to investigate CNS leukemia involvement in all leukemia patients. None of the patients underwent any additional procedures for study purposes. 1 ml of CSF was removed during the routine procedure. The identity information of the patients is kept confidential.

### SAMPLE GROUPING

Two groups: a control group (N=21) and a positive group (N=5). CSF leukemia is diagnosed by cytopathology. The positive group indicated that the patients tested positive for leukemia. The control group tested negative to the leukemic cells.

### SAMPLE COLLECTION AND TRANSPORT

The samples were transported via dry ice to the lab for FTIR spectroscopy. The samples were collected in batches with the patient’s numbers tagged along samples that were collected.

### SAMPLE PREPARATION AND FTIR ANALYSIS

The machine was cleaned with 70% isopropyl and distilled water before taking a background spectrum. A background spectrum was first taken before the samples were analyzed. The samples were vortexed gently to prevent cell death. Fluid was taken from each of the samples using 1 microliters pipette and dropped on the diamond crystals on the FTIR machine. They were gently dried under nitrogen gas for 3 minutes each and for each patient three replicas were made. As a result, all samples were dried for 3 minutes for the FTIR investigations. The reason for a gentle drying with nitrogen gas is the removal of free water in the feeder environment. The time for drying was chosen according to the optimization studies with different drying minutes and 3 minutes was chosen for the best results. Following the drying process, the spectrum was then started to be recorded in the 4000-850 cm-1 wave count range, 4 cm-1 resolutions, and 128 scan number. The spectrum data was recorded and analyzed using Opus Software (version 8.5, Bruker Optic, GmbH). From each sample 3 replicates as 1 microliter for each were used to record the spectrum. The average of these three spectra was then used to examine that sample’s molecular properties. For the first derivative the following was calculated: amide I band, amide II band, CH2, COO symmetric stretching, Phosphate band, Glucose band and DNA band and intensity ratio were measured).

### ANALYSIS OF FTIR DATA

Prior to performing the spectrum analysis, a full spectral baseline correction was performed, which is frequently done to eliminate a sloping or curved baseline. The middle 80% of peak heights in the spectral data were used to calculate band locations (wavenumber values). Some of the bands were not that clearly visible in the first derivative so the band intensity instead of band area from second derivative was used for the calculation. After integrating the band area values using the area beneath the target band as a guide from first derivative or using vector normalization to measure the band’s intensity from second derivative, ratios of band areas or band intensities were also calculated. We also considered the half-wavenumber of the CH_2_-antisymmetric band when calculating the half bandwidth. Each result presented here is the mean of three independent samples. The average spectra were normalized with respect to the relevant bands in numerous images scattered throughout the thesis to graphically depict the differences between the groups.

### STATISTICAL ANALYSIS

Each indicator of each sample measures the coefficient of variation (standard deviation/average), summarizes the data as the average and standard deviation of each group. GraphPad (version 9.0) was used to analyze statistics. A one-way ANOVA was used to evaluate the band area/intensity ratio based on normality test results, while the Kruskall Wallis test was used to analyze the wave number and half-band width values. Means and standard deviations (SDs) were used to represent the data. The P value is less than 0.05 () or less than 0.05 (=) and is used to determine the statistical significance (*, +p 0.05; **, +++p 0.01; ***, +++p0.001).

## 3. Results

The application of FTIR spectroscopy revealed distinct spectral differences in the CSF of patients with CNS involvement, characterized by specific molecular vibrations that correlate with pathological changes. These results were compared with those obtained from traditional methods, highlighting the enhanced sensitivity and specificity of FTIR spectroscopy in identifying critical biomarkers.

In this research, detection of CNS involvement in acute leukemia was performed using ATR-FTIR Spectroscopy.

### 4.1 RESULTS OF FTIR SPECTROSCOPY STUDIES

The band assignment for the representative spectrum of control patient CSF sample is displayed in Table 4.1 which is also labeled in Figure 4.1 The second derivatives’ spectra was only used for the carbonyl ester triglyceride and olefinic bands. To obtain data on DNA concentration, aliphatic chain length, protein structural and conformational changes, protein concentration, saturated and unsaturated lipid concentration in addition to carbonyl, carbohydrate, and nucleic acid contents, the area and intensity ratios of the studied bands were calculated. While the other parameters were evaluated using band area ratios, the olefinic, saturated lipid, carbonyl ester, and carbohydrate contents were computed using band intensity ratios from the second derivative spectra.

**Table 4.1:**
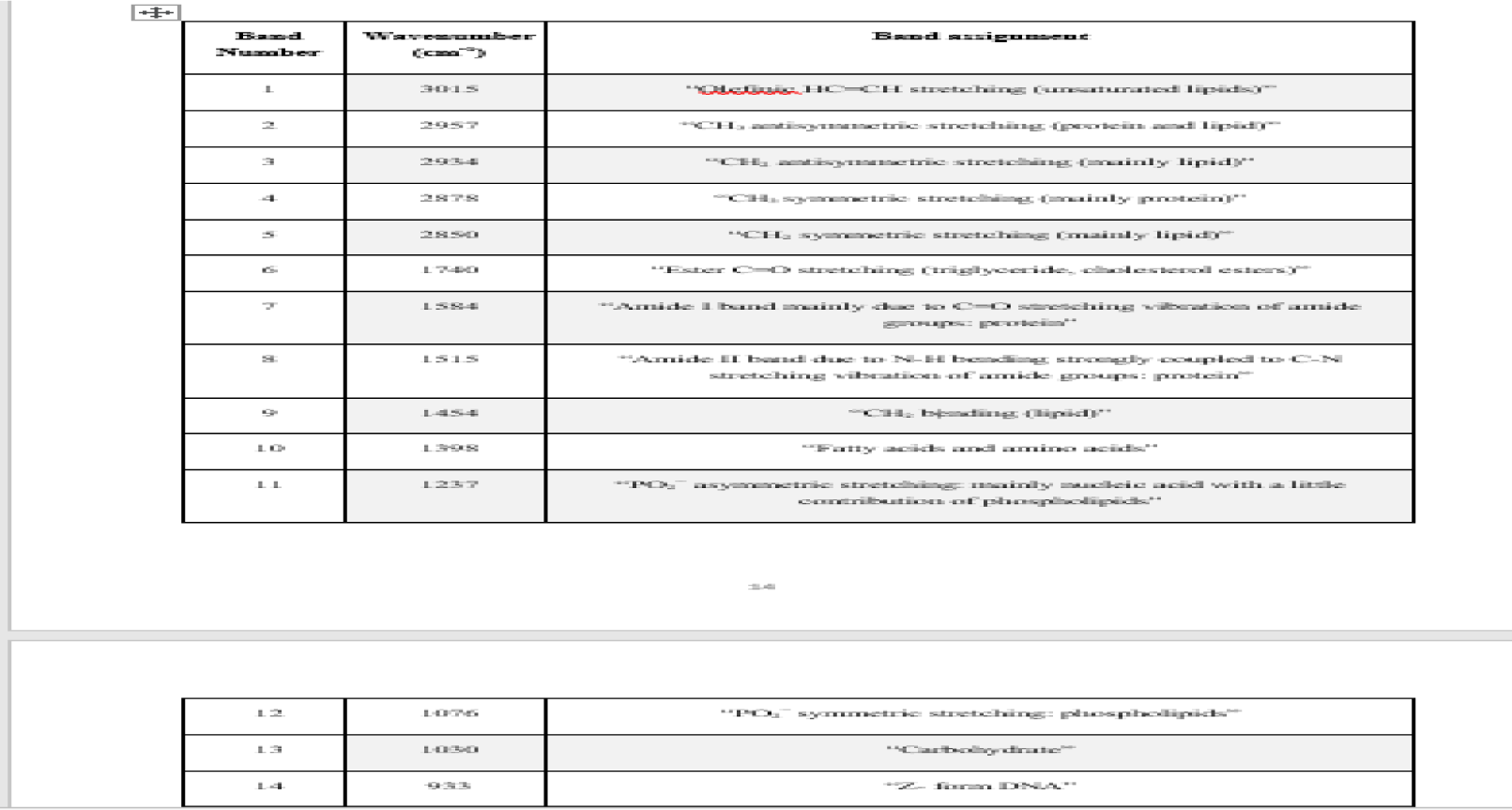
The Band Assignment of An IR Spectrum Between 3000-750 Cm-^1^ Wavenumber Region. (Algrubi Et Al., 2022; Dogan Et Al., 2021; Ustaoglu Et Al., 2021; 2018; Yonar Et Al., 2018; Garip Et Al.,2007;2009).

The alterations in lipid composition and structure were studied using the band area/intensity ratios of a few different lipid functional groups in the C-H stretching region (3015-2800 cm-1), as well as the C=O stretching vibrations of carbonyl ester groups. The calculated ratios are listed below: olefinic HC=CH stretching/ (CH2 antisymmetric stretching + CH2 sym stretching) as unsaturated lipid content, Ester C=O stretching/ (CH2 antisymmetric stretching + CH2 symmetric stretching) as carbonyl content; CH2 antisym stretching/ (CH2 antisym stretching + CH2 sym stretching) as saturated lipid content; and CH2 antisymmetric stretching/CH3 antisymmetric stretching as acyl chain length.

**Figure 4.1:**
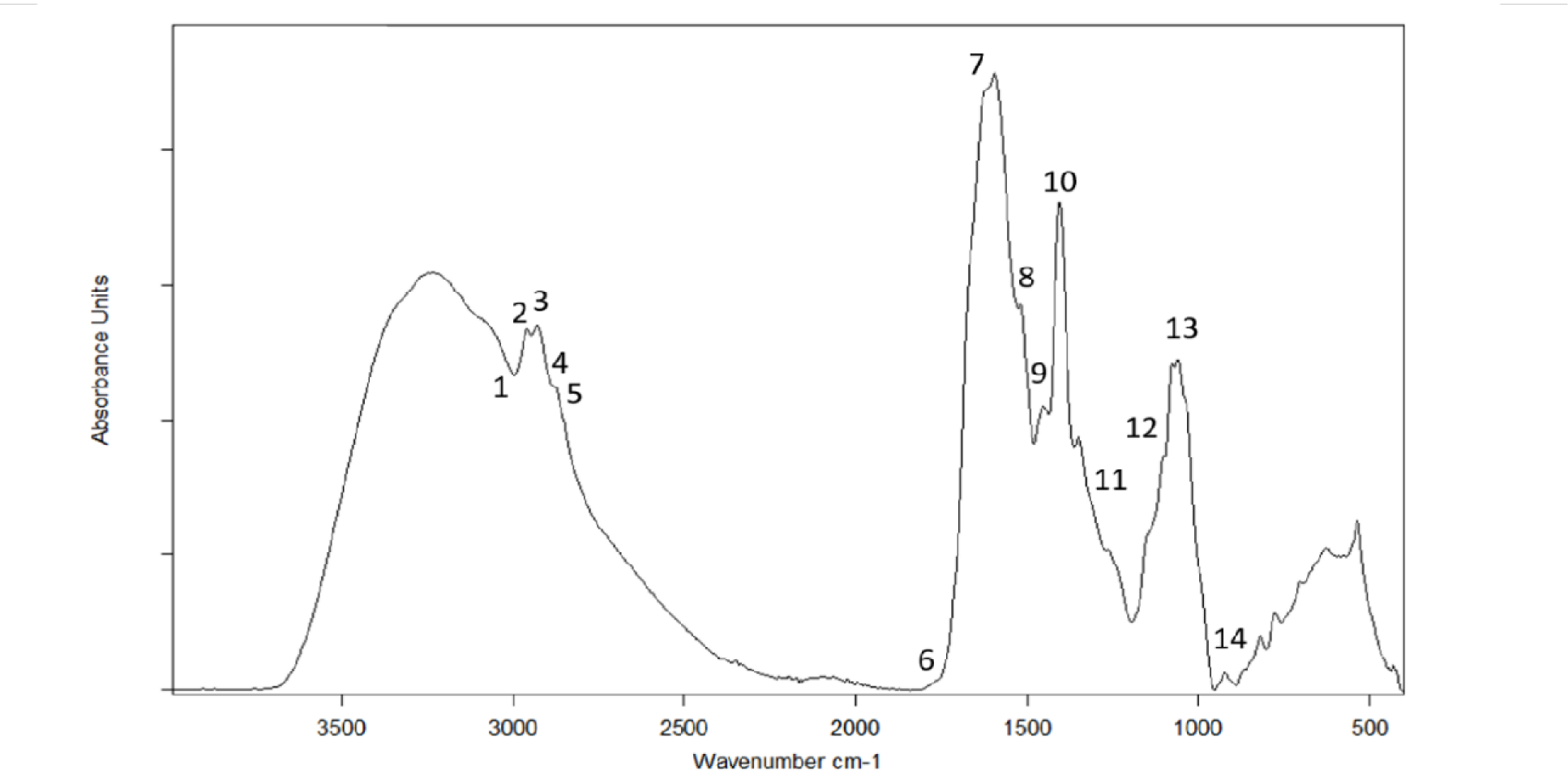
The Representative Spectrum of CSF Sample of Control Patient Between 4000-400 Cm^1^ Wavenumber Region.

**Figure 4.2:**
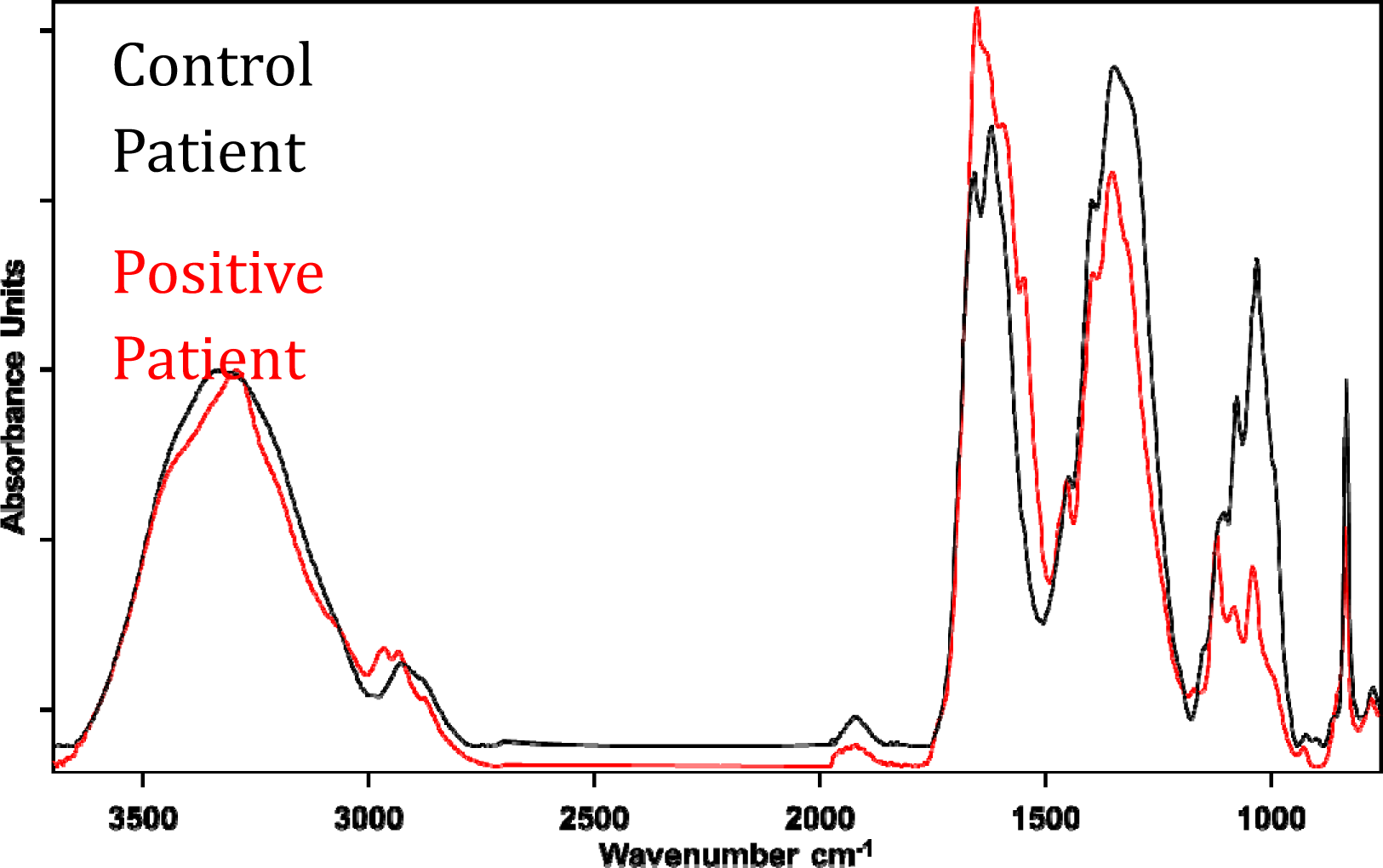
Representative IR spectra of the control and positive patients’ CSF samples: (A) absorbance spectra in the 3750–900 cm^−1^ region (normalized with respect to the amide A band located at 3330 cm^−1^), (B) second derivative vector normalized spectra in the C-H region (3050–2800 cm^−1^).

**Table 4.2:** Changes in the band intensity of olefinic band at 3012 cm ^1^, the band area/intensity ratios of various functional groups and the wavenumber and bandwidth values of CH_2_ antisymmetric stretching bands of control (C) and positive patient (PP) groups.

| Band Intensity | Explanation of the calculations | Control (n=21) | Positive Patient (n=5) |
| --- | --- | --- | --- |
| Olefinic | Olefinic Content | 0.007 ± 0.005 | 0.004 ± 0.011 |
| **Band Area/Intensity Ratios** |  |  |  |
| Olefinic / $\text{CH}_2$ antisym + $\text{CH}_2$ sym | Total unsaturated lipid content | 0.287 ± 0.293 | 0.042 ± 0.046<br>**** |
| $\text{CH}_2$ antisymmetric / $\text{CH}_2$ antisymmetric + | Total saturated | 0.684 ± 0.682 | 0.140 ± 0.114 |
| CH<sub>2</sub> sym | lipid content |  | **** |
| Ester C=O stretching/<br>CH<sub>2</sub> antisymmetric +<br>CH<sub>2</sub> sym | Carbonyl<br>content | 1.430 ± 0.281 | 1.77 ± 0.177 |
| CH<sub>2</sub> antisym / CH<sub>3</sub><br>antisym | Lipid acyl<br>chain length | 2870 ±<br>1.94 | 1699.48 ±<br>45.240 |
| Amide 1 / Amide 1 +<br>Amide 2 Int | Total protein<br>content | 0.620 ±<br>0.620 | 0.653±<br>0.689 |
| Amide I / Amide II | Protein<br>conformation<br>al change | 1.672 ±<br>1.680 | 1.998 ±<br>2.263 |
| Phosphate sym /<br>Amide I band area. | Nucleic acid<br>content | 0.362 ±<br>0.344 | 0.101 ±<br>0.100<br>**** |
| DNA band /<br>Phosphate sym band<br>intensity | DNA content | 0.737 ±<br>0.728 | 0.836 ±<br>0.823 |
| RNA band /<br>Phosphate sym band<br>intensity | RNA content | 539.48 ±<br>152.50 | 0.564 ±<br>0.157<br>*** |
| Glucose Area/Amide<br>1 Area | Glucose<br>content | 0.403 ±<br>0.405 | 0.460 ±<br>0.487 |
| **Wavenumber Value** |  |  |  |
| CH<sub>2</sub> antisymmetric<br>stretching | Lipid order | 2872.8 ± 59.2 | 2742.8 ±<br>212.491 |
| Amide I |  | 1.855 ± 0.968 | 1726 ± 55.34 |
| **Bandwidth Value** |  |  |  |
| CH<sub>2</sub> antisymmetric stretching | Lipid dynamic | 2872.8 ± 59.2 | 2742.8 ± 212.491 |

*The degree of significance was denoted as $p < 0.05^*$ , $p < 0.01^{**}$ , $p < 0.001^{***}$*

The signal intensity or to be more precise, the area of the absorption bands in the FTIR spectrum provides information about the relative concentrations of the associated functional groups (Baloglu et al., 2015). Differences in cross-sectional thickness can lead to concentration-dependent spectral variations in absorbance values. Therefore, band area ratios were used instead of band area alone to calculate contextual and structural parameters (Ustaoglu et al., 2021).

Total lipid amount of the system was calculated by taking the band intensity of CH_2_ antisymmetric to the sum of the intensity of both CH_2_ antisymmetric and symmetric stretching bands. Total unsaturated lipid content was calculated by taking the band intensity ratio of olefinic band to the sum of the intensity of both CH_2_ antisymmetric and symmetric stretching bands. Moreover, by checking the band intensity of olefinic band itself, concentration information about the unsaturated lipids in the system can be obtained. As seen from Figure 4.3, there is a significant decrease in both total unsaturated and saturated content of positive patient group compared to the control group (Table 4.2).

The carbonyl content, otherwise known as the triglyceride content of the system was derived by taking ratios of the intensity of Ester C=O stretching band to the sum of CH_2_ antisymmetric and CH_2_ symmetric stretching bands. There was a noticeable decrease in the positive patients relatively compared to the control groups.

The nucleic acid content was derived by obtaining the band area ratio of the sum of PO2– symmetric and Amide I area. The nucleic acid content of the control group was increased as seen in figure 4.7. even though the decrease was not significant, this increase could be correlated with the decrease in RNA content and DNA content. Which are represented in figure 4.8 and 4.9 respectively Total protein concentration was calculated by taking the ratio of the band area of Amide I to Amide I and Amide II intensity bands. As seen from Fig. 4.6 alongside with Table 4.3 the protein content was notably increased in the positive patient when compared to the control groups the protein content was not significant when comparing to the control groups while taking the value closest to the control group (non-significant compared to the control).

### TOTAL SATURATED AND UNSATURATED LIPID CONTENT

**Figure 4.3:**
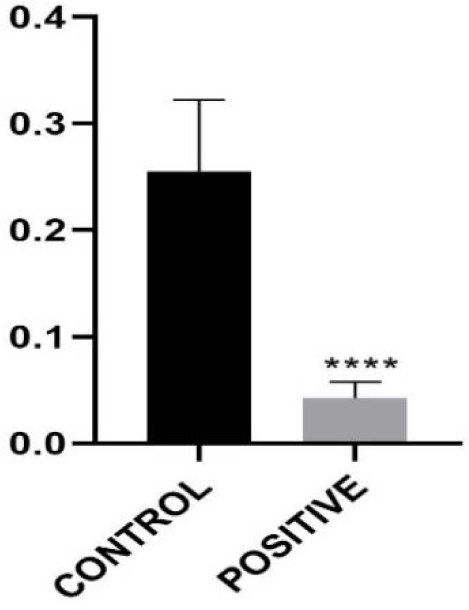
Olefinic HC=CH Stretching / CH2 Anti Symmetric + CH2 Sym Intensity.

**Figure 4.4:**
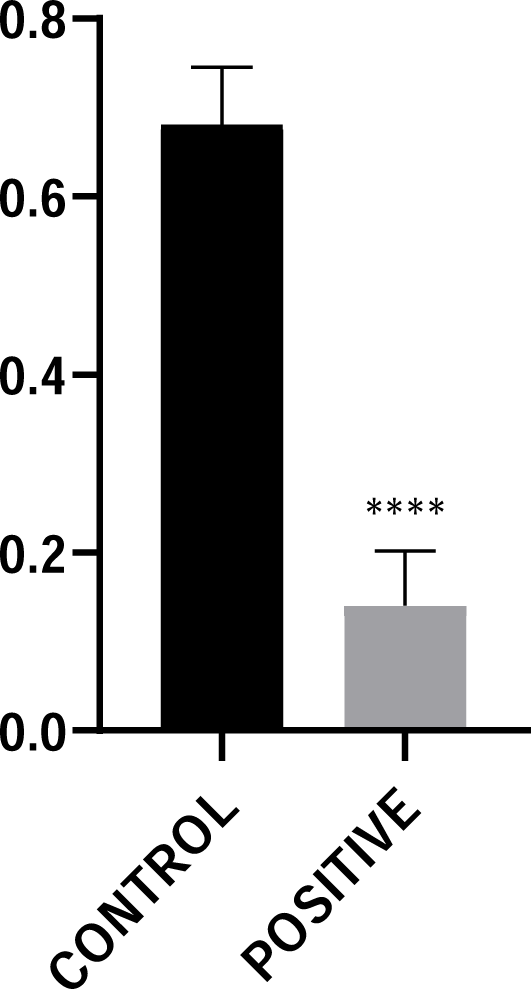
CH_2_ anti sym/ CH2 anti sym + CH2 sym intensity.

**Figure 4.5:**
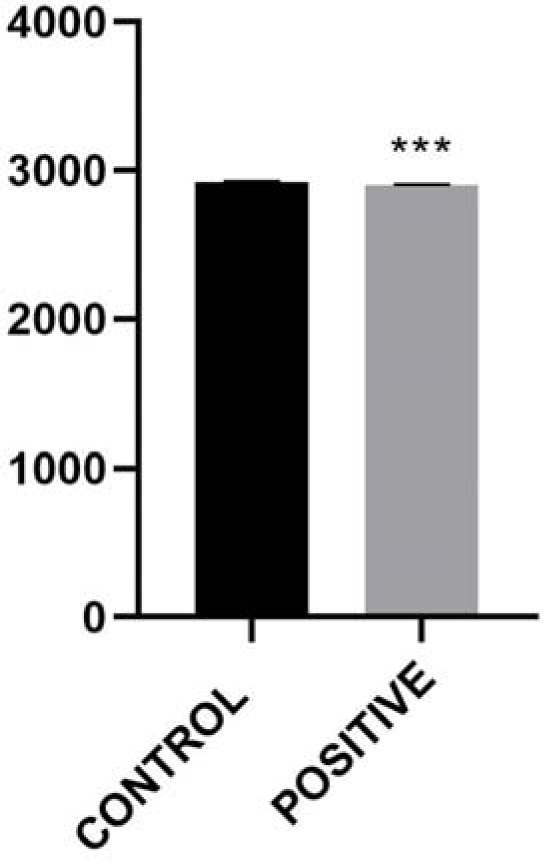
CH_2_ anti symmetric stretching band wavenumber.

**Figure 4.6:**
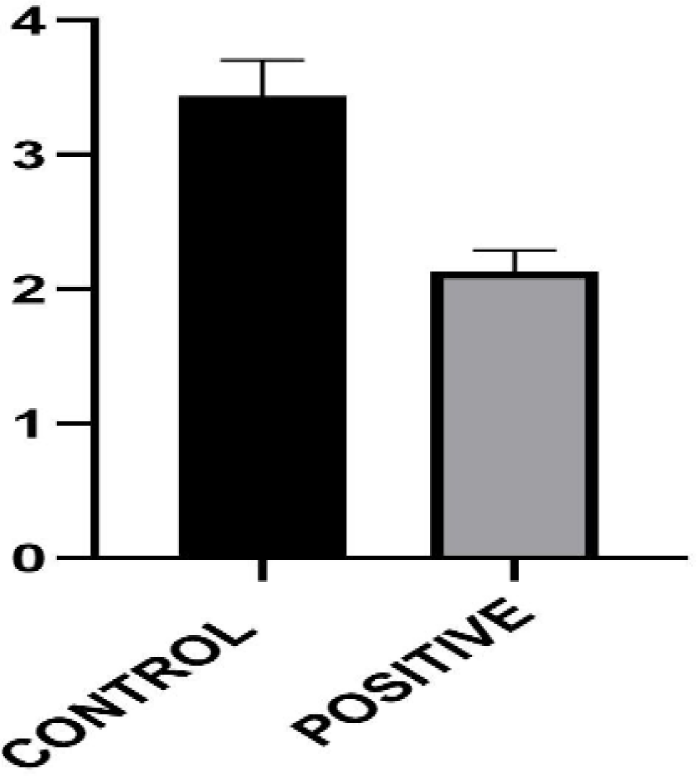
CH_2_ anti symmetric stretching band half-bandwidth.

### LIPID ORDER / DISORDER AND LIPID DYNAMICS

Moreover, the variations in the chain lengths of the lipids was also calculated by dividing CH2 antisym stretching band area to CH3 antisym stretching (at 2958 cm-1) band area (Algburi et al., 2022). Lipid acyl chain length of positive patient group was significantly lower than control patient group (Figure 4.4, Table 4.2).

The shifts in the wavenumber of the CH2 antisymmetric stretching band give information about a structural parameter called lipid order. Lipid order represents the flexibility of the lipid acyl chains. As seen from Figure 4.5 and Table 4.2, there was a change in the wavenumber of CH2 antisym band to lesser wavenumber values in positive patient group compared to the control implying an increase in lipid order. Moreover, variations in the bandwidth of the same band give information about lipid dynamics as a functional parameter. There was a significant increase in the bandwidth of CH2 antisym band in the positive patient group compared to the control (Fig 4.6, Table 4.2) indicating an increase in lipid dynamics.

Figure 4.3 should include olefinic intensity, total unsaturated, saturated content, lipid acyl chain length, CH_2_ wavenumber and CH_2_ bandwidth.

### 4.3 NUCLEIC ACID CONTENT AND DNA CONTENT

**Figure 4.7:**
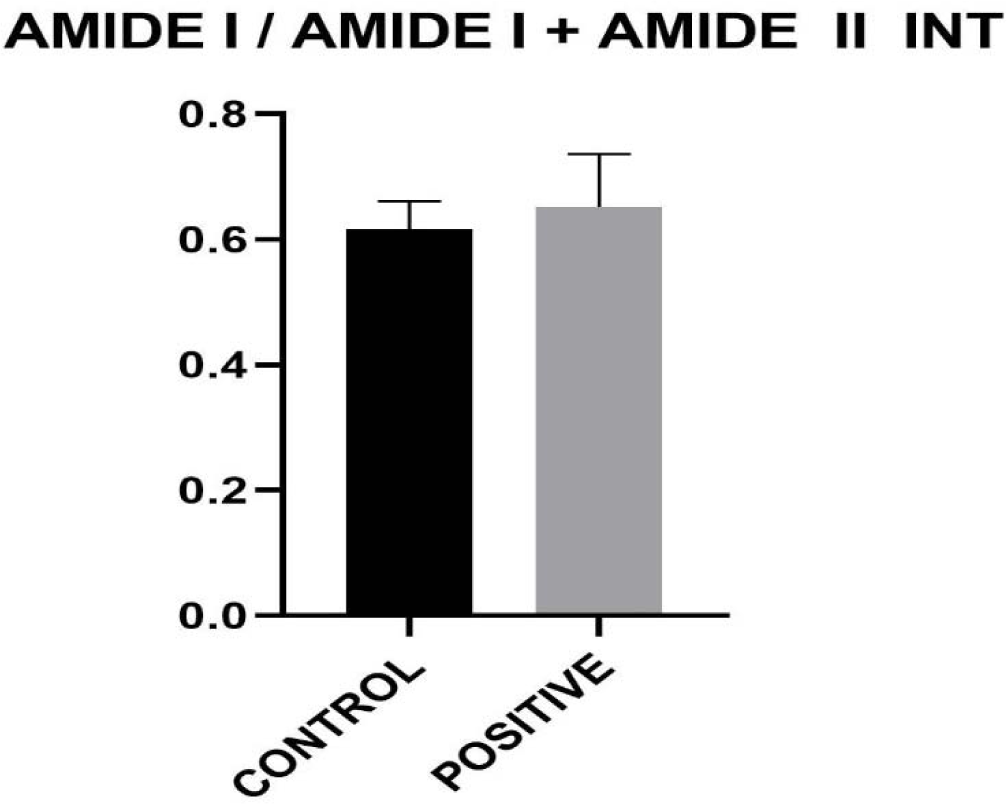
Amide I / Amide I and Amide II intensity

**Figure 4.8:**
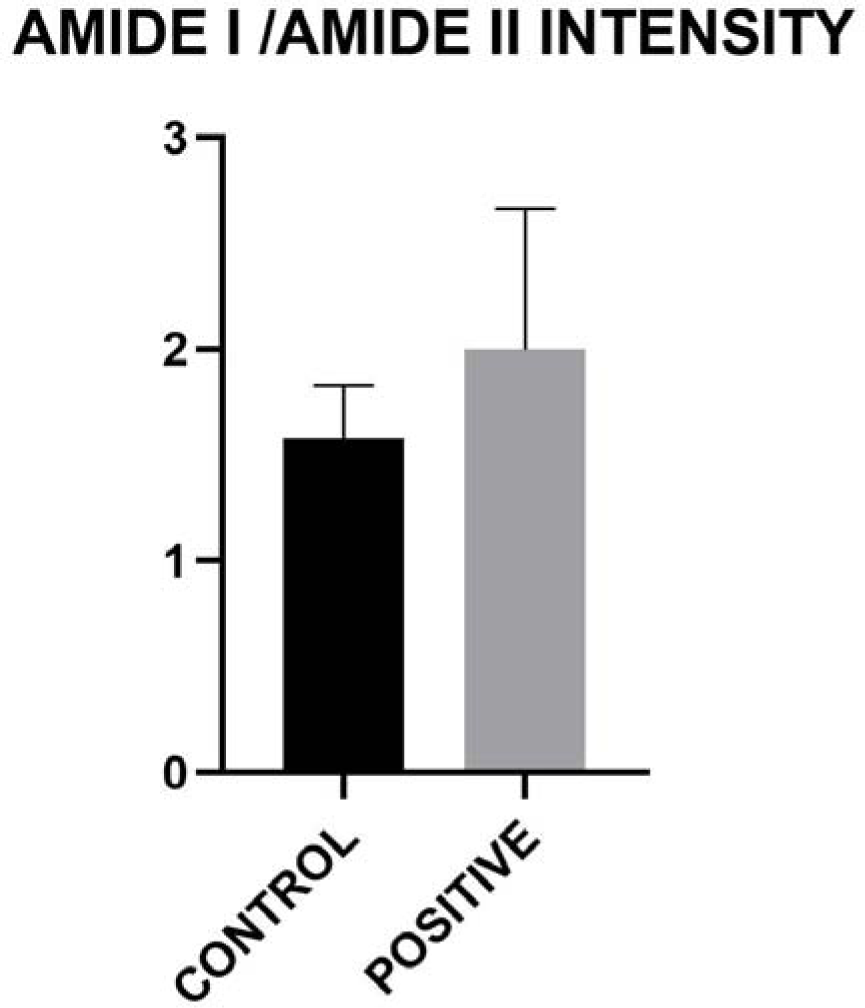
Amide I/ Amide II intensity.

**Figure 4.9:**
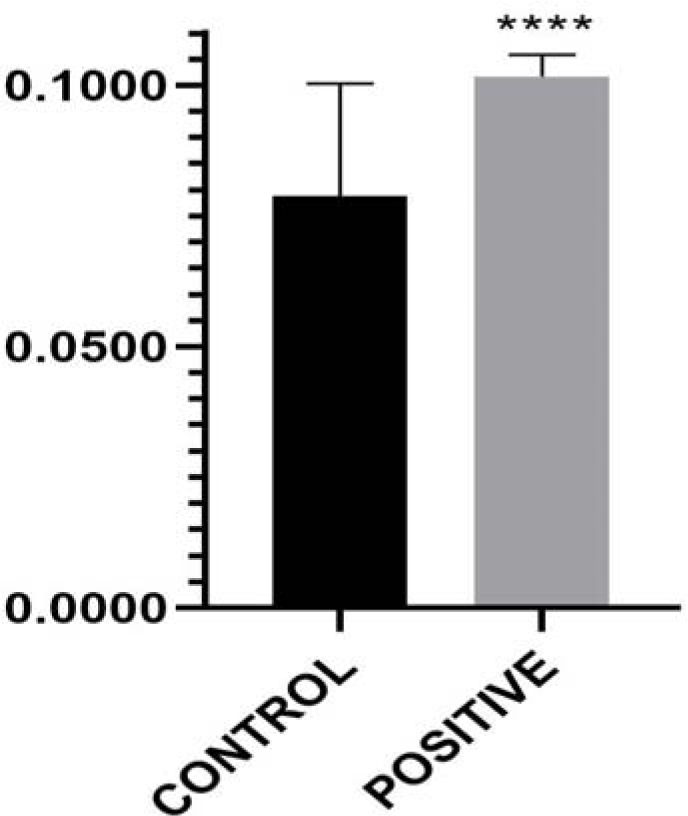
Phosphate sym / Amide I band area.

**Figure 4.10:**
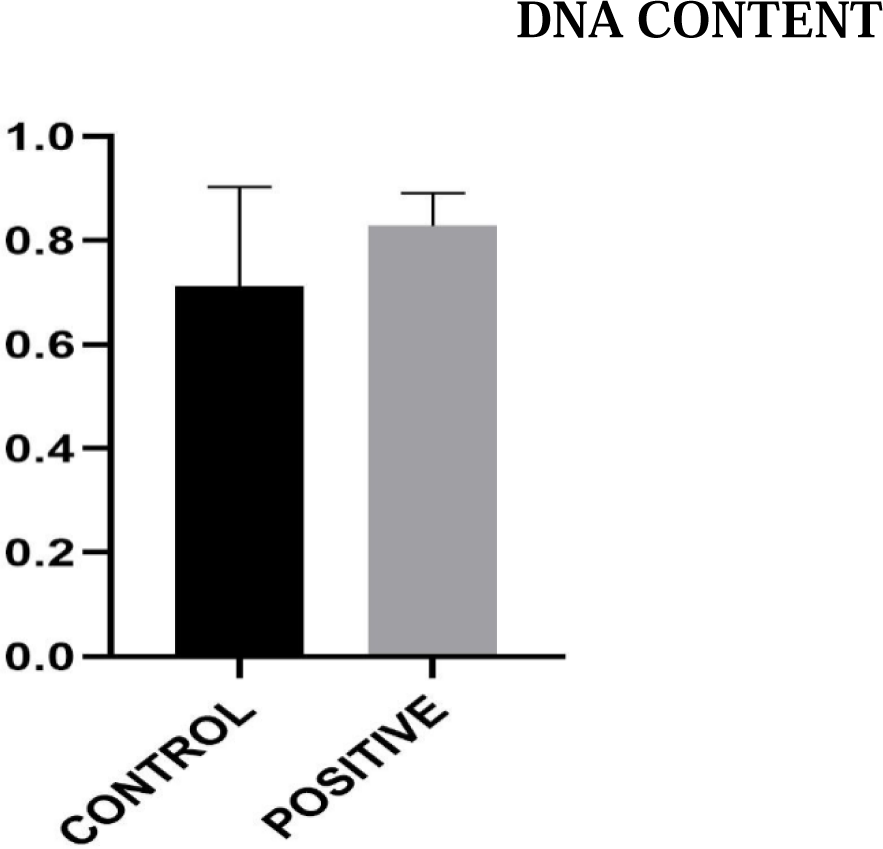
DNA band / Phosphate sym band intensity.

**Figure 4.11:**
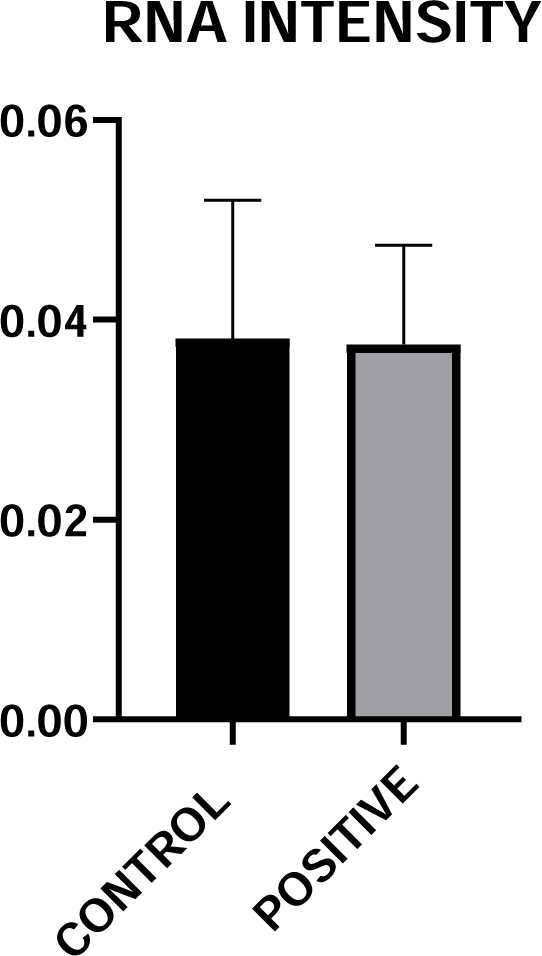
RNA band / Phosphate sym band intensity.

**Figure 4.12:**
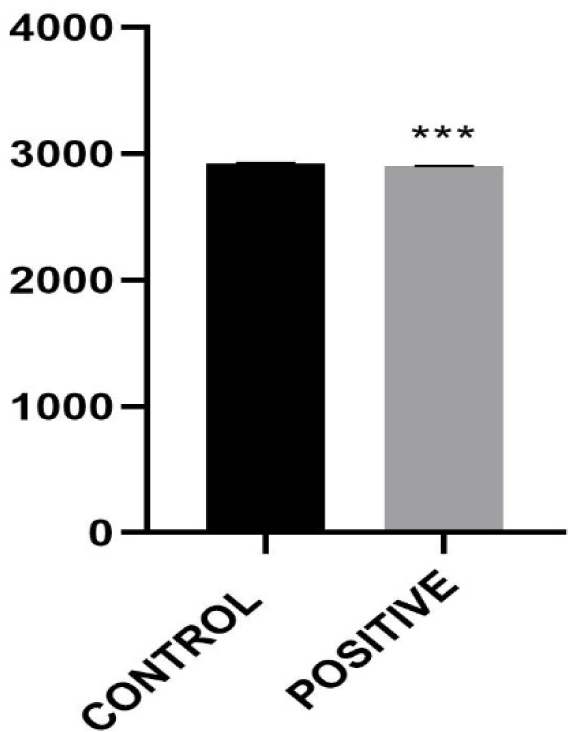
Olefinic HC=CH stretching / CH_2_ anti sym + CH_2_ sym intensity.

**Figure 4.13:**
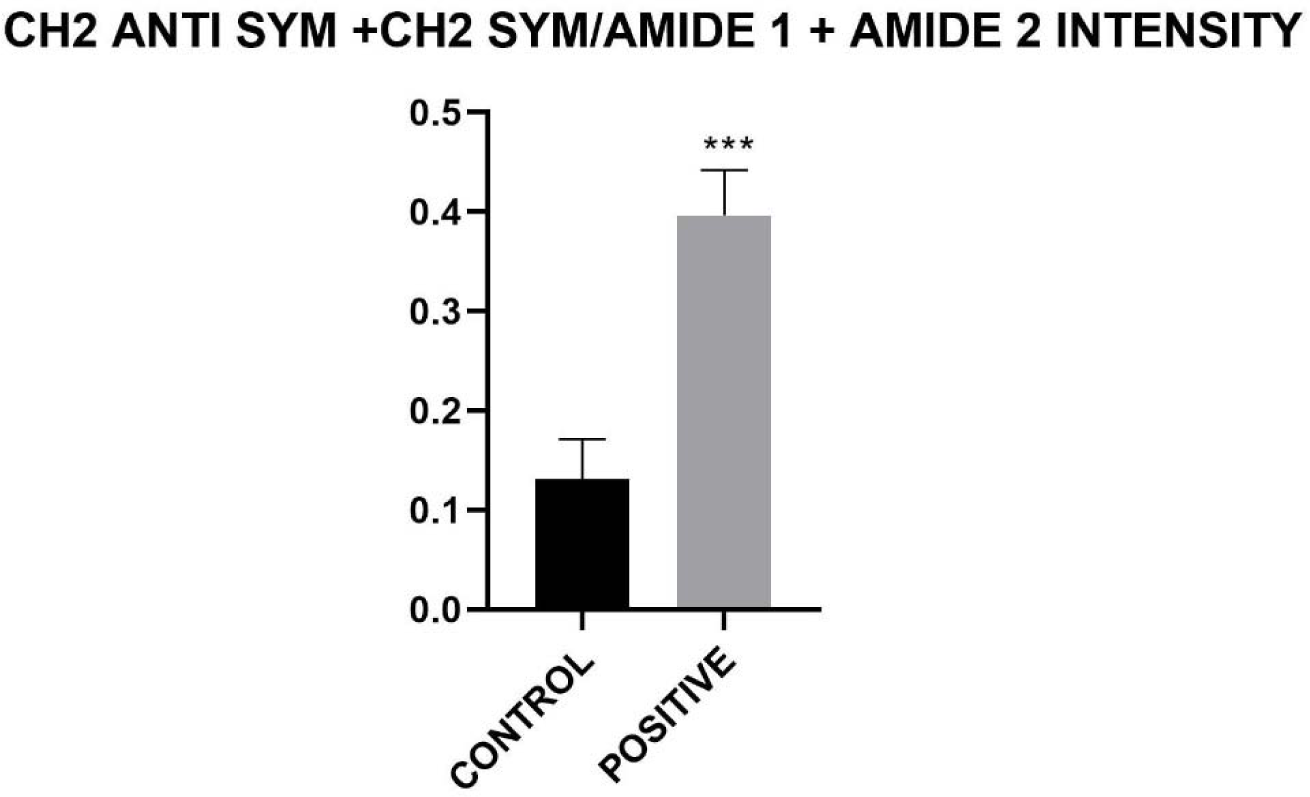
CH_2_ Anti sym and CH_2_ sym/Amide 1 and Amide II intensity.

**Figure 4.14:**
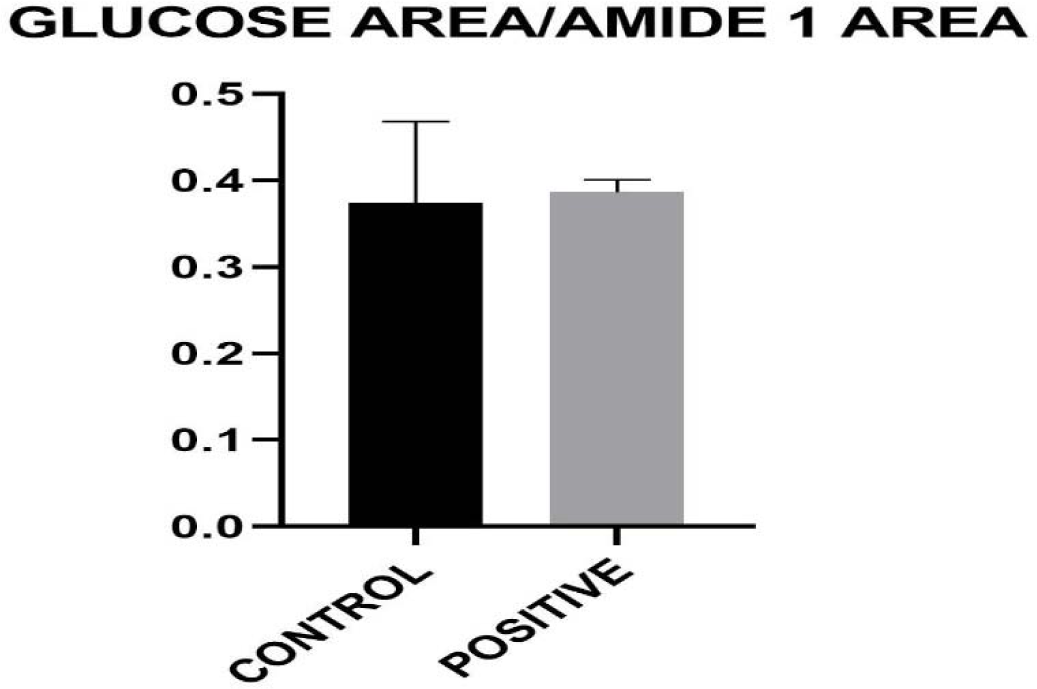
Glucose area/ amide I area.

**Figure 4.15:**
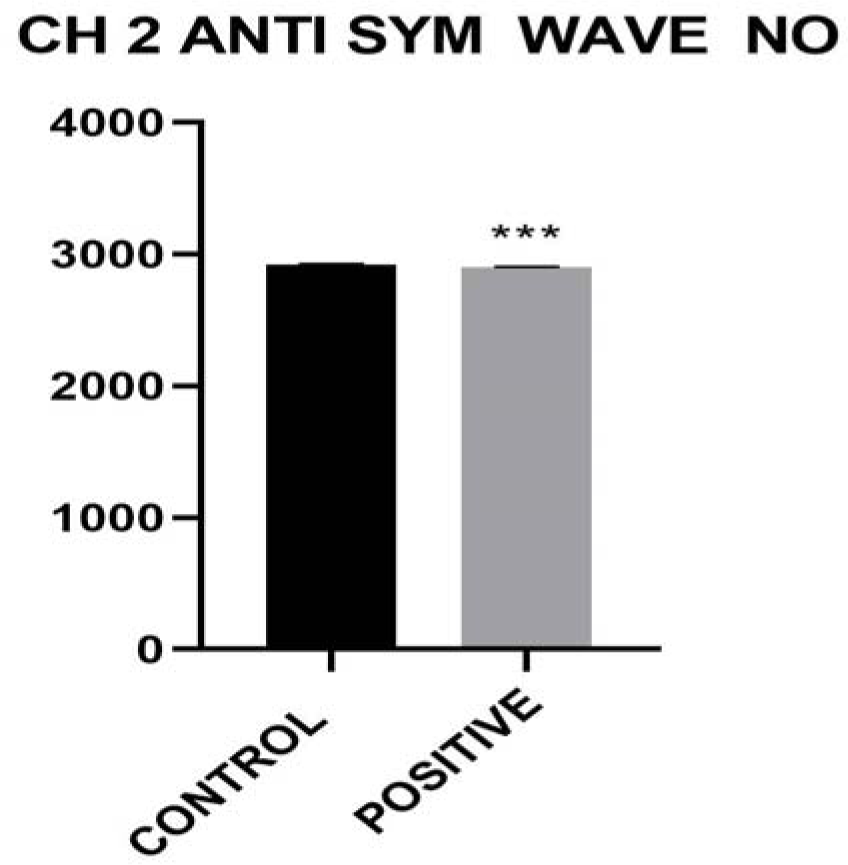
CH_2_ anti symmetrical wave number.

**Figure 4.16:**
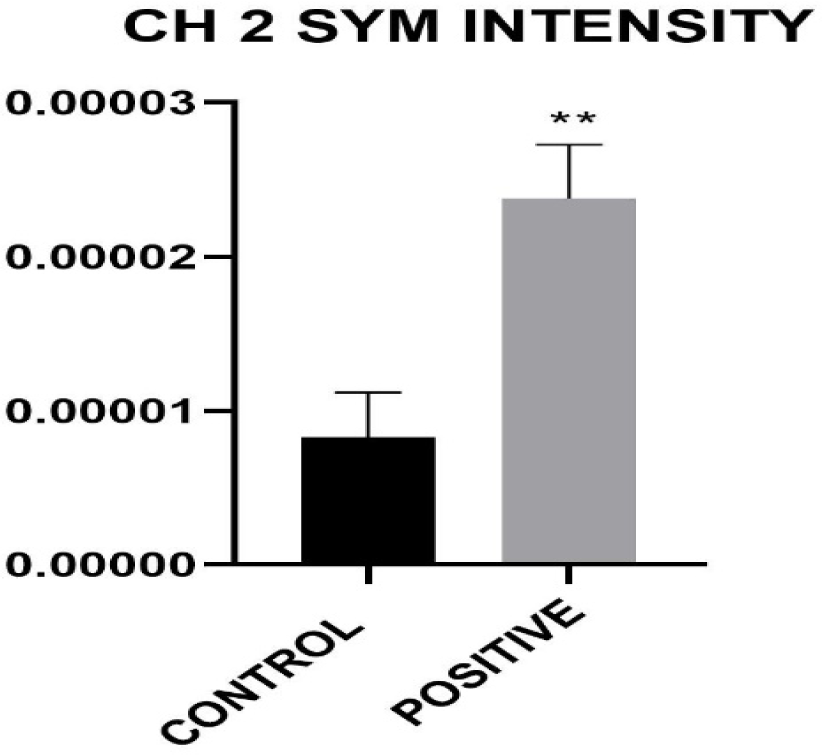
Olefenic intensity

## 4. Discussion

The findings from this study underscore the potential of FTIR spectroscopy as a superior diagnostic tool for early detection of CNS involvement in leukemia. The ability of FTIR to provide detailed molecular insights offers significant advantages over conventional methods, potentially leading to earlier intervention and improved patient outcomes. This discussion explores the implications of these findings for clinical practice and future research.

Pediatric leukemia, the most frequent childhood cancer in children worldwide, comes in two main varieties: acute lymphocytic leukemia (ALL) and acute myelogenous leukemia (AML). Patients with leukemia experience abnormal white blood cell development, and stem cells grow into lymphoblasts (leukemia cells) Samantha et al. (2017). The possibility to detect childhood leukemia early using merely an investigation of the serum FTIR spectra appears to be a promising tool for everyday medical practice. Results from the present research work has indicated that the ratios of peak heights could differentiate the serum of leukemia patients from that of healthy controls., which demonstrated the greatest differences in their peak regions it was observed that the plasma spectra of healthy children were noticeably different from those of patients with chronic lymphocytic leukemia in terms of their peaks differences from the Amide, lipids, nucleic acid and glucose bands. Additionally, analysis of the observed spectra at those specific regions yielded a precise classification of the healthy and positive samples that perfectly matches with clinical information” (Erukhimovitch et al. 2006: Chaber et al., 2021).

The 2nd derivative spectra of control and positive patients in 3050-2800 cm-1 wavenumber region shows the decrease in both saturated and unsaturated lipid content in posisitve patient group in figure 4.3 The total saturated and unsaturated lipid content in figure 4 and 5 are represented by the CH2 anti sym/ CH2 anti sym + CH2 sym intensity and Olefinic HC=CH stretching / CH2 anti sym + CH2 sym intensity respectively. They were taken from the ssecond derivative due to This shows a significant decrease in the positive patients when compared to the control group. There is chronic oxidative stress in patients with leukemia (Kruk et al., 2017), this leads to decrease in total lipid content and increase in lipid peroxidation. The subsequent peroxidation attacks the double bonds which further decreases the reaction oxygen specie, not only is the ROS attacked but also the membrane proteins are affected,. This can therefore be attributed to decrease, subsequently, the lipid order / disorder and lipid dynamics represented by CH2 anti symmetric stretching band wavenumber and CH2 anti symmetric stretching band half-bandwidth in figure 6 and 7 shows a significant difference between the two groups. There is a decrease in wavenumber and bandwidth respectively. This could be further attributed to the increase in the lipid order due to increase in lipid peroxidation. Hence, there is a corresponding decrease in the fluidity of the system in these positive patients. The nucleic acid content and DNA content represented in figure 8 and 9 showed a significant increase in the positive patients, this increase could be correlated to the increase in the DNA and RNA content. There is rapid division of the cells in the positive patient, hence replication is doubled.

The glucose content of the system is not significantly increased when comparing the control to the positive groups. The tumor cells cause high level production of IGFBP1 from the adipose tissue to facilitate insulin sensitivity in the positive patients. Additionally, obesity/diabetes associated chronic inflammation also act to promote the spread and survival of tumors (Deng et al., 2016).

## 5. Conclusion

FTIR spectroscopy represents a significant advancement in the diagnostic evaluation of pediatric leukemia patients with potential CNS involvement. The research shows that FTIR spectroscopy can differentiate between the serum and plasma spectra of leukemia patients and healthy individuals based on peak ratios at specific regions. The findings are consistent across different studies and show significant differences in peak intensities related to carbohydrates, amino acids, and proteins. The use of FTIR spectroscopy for early diagnosis can greatly improve the chances of successful treatment, and further research in this area is warranted. Acute lymphocytic leukemia (ALL) and Acute myelogenous leukemia (AML) are the two primary subtypes of pediatric leukemia, the most common cancer in children globally. Intriguing differences between the FTIR spectral signatures of leukemic and healthy serum were found, according to the results. These distinctions might offer a practical method for FTIR spectroscopy-based early ALL diagnosis in children, reducing the need for invasive procedures and accelerating patient diagnosis.

## Notes

### Competing Interest Statement

The authors have declared no competing interest.

